# Chromosome compartment assembly is essential for subtelomeric gene silencing in trypanosomes

**DOI:** 10.1101/2025.02.21.639481

**Authors:** Luiza Berenguer Antunes, Tony Isebe, Oksana Kutova, Igor Cestari

## Abstract

Genome three-dimensional organization is essential for the coordination of eukaryote gene expression. The chromosomes of the pathogen *Trypanosoma brucei* contain hundreds of silent variant surface glycoprotein (VSGs) genes in subtelomeric regions. However, *T. brucei* transcribes a single VSG gene and periodically changes the VSG expressed by transcriptional or recombination mechanisms, altering its surface coat to escape host antibodies by antigenic variation. We show that VSG-rich silent subtelomeric regions form distinct chromosome compartments from transcribed regions, with subtelomeric compartments of different chromosomes co-interacting. We uncovered chromatin-associating factors at the boundaries of transcribed and silent compartments. Among these, repressor activator-protein 1 (RAP1) marks the compartment boundaries and spreads over silent regions. Inactivation of phosphatidylinositol phosphate 5-phosphatase removed RAP1 from compartment boundaries and subtelomeric regions, disrupting compartment assembly and derepressing all VSG genes. The data show spatial segregation of repressed from transcribed chromatin and phosphoinositides regulation of silent compartment assembly and genome organization.

## Introduction

Eukaryote chromosomes are typically organized into megabase (Mb) scale compartments defined as transcribed A compartment (euchromatin) and silent B compartment (heterochromatin) ^1,2^. Compartments are further organized into topologically associating domains (TADs), sub-TADs, and loops ^2,3^. This spatial organization provides levels of compaction and coordination of gene expression by placing spatially distal genomic regions in proximity, such as genes and their promoters and enhancers, or spatially parting transcribed from repressed regions ^4–6^. The megabase (Mb)-long diploid chromosomes of the protozoan pathogen *Trypanosoma brucei* are organized into core regions containing polycistronic units (PTUs) encoding housekeeping genes, subtelomeric regions containing largely silent variant surface glycoprotein (VSGs) genes, and in some chromosomes, telomere-proximal expression sites (ESs) from where VSG genes can be transcribed ^7^. The PTUs contain dozens of genes lacking canonical RNA polymerase II promoters and are co-transcribed into polycistronic RNAs processed by trans-splicing ^8^.

*T. brucei* infect humans and animals, causing sleeping sickness. These parasites express a homogeneous surface coat composed of ~ 10^7^ VSG proteins ^9,10^, which they periodically switch to evade host antibody clearance by antigenic variation. VSG proteins contain a variable surface exposed N-terminal domain, and a conserved C-terminal region attached to the cell’s surface by a glycosylphosphatidylinositol anchor ^11,12^. There are over 2,500 VSG genes and pseudogenes in this organism genome, found primarily in subtelomeric regions which function as a storage of VSG sequences ^7^. However, a single VSG gene is transcribed at a time from one of the 20 ESs. The change in VSG expressed occurs by transcriptional switching among ESs or by recombination of VSG genes within the active ES, altering the parasite surface coat. The mechanisms regulating VSG monogenic expression remain poorly understood. The active VSG is transcribed from a nuclear site termed ES body (ESB) ^13^. Specific factors, such as ESB-1 and VSG exclusion 1 and 2, associate with the active ES in the ESB and are required for VSG gene transcription ^14–16^. In contrast, the silencing of the remaining ESs depends on repressor-activator protein 1 (RAP1) ^17^, which binds DNA sequences flanking the ES VSG genes ^18,19^. RAP1 silencing function is controlled by a phosphoinositide regulatory system, disruption of which results in VSG derepression and switching ^19,20^. Specifically, PI(3,4,5)P3 is an allosteric regulator of RAP1, and its nuclear accumulation displaces RAP1 from ESs resulting in VSG transcription ^18,19^. A nuclear phosphatidylinositol phosphate 5-phosphatase (PIP5Pase) dephosphorylates PI(3,4,5)P3 and maintains ES repression. Its catalytic inactivation results in ES transcription and VSG switching ^19^, indicating that PIP5Pase plays a role in ES chromatin organization.

*T. brucei* chronic infection relies on the vast repertoire of subtelomeric VSG genes to escape host antibodies. The parasites’ inability to maintain their repression results in multiple VSGs expressed ^18,19^ and their immune clearance during infection ^21^. It remains unknown how hundreds of subtelomeric VSG genes from all 11 Mb chromosomes are repressed. Here, we show that silencing subtelomeric VSG genes in *T. brucei* entails spatial chromatin organization. We show that chromosomes are organized into compartments, TADs, sub-TADs and loops, with subtelomeric regions forming distinct silent compartments from core transcribed regions and interacting among chromosomes. Using cross-linking mass spectrometry, we identified chromatin-binding proteins associated with compartment boundaries, including RAP1, which spreads over silent subtelomeric compartments. The knockdown of PIP5Pase disrupts chromosome compartment contacts and displaces RAP1 from subtelomeric regions, resulting in the transcription of all subtelomeric VSG genes. The data indicate that silencing subtelomeric genes entails spatial chromosome organization into chromatin compartments and is PIP5Pase and RAP1 dependent. It also shows a role for phosphoinositides in controlling chromatin spatial organization.

## Results

### Spatial chromatin protein cross-link network at ~30 Å resolution

To identify chromatin regulatory proteins, we performed *in vivo* chemical cross-linking and immunoprecipitation of PIP5Pase followed by mass spectrometry (XLMS). We used a *T. brucei* bloodstream form line expressing an endogenously tagged PIP5Pase with three C-terminal V5 epitopes (42 amino acids), which we showed does not interfere with enzyme activity, interactions, or localization ^18–20^. We cross-linked cells with disuccinimidyl suberate (DSS), a cell-permeable non-cleavable cross-linker with amine-reactive N-hydroxysuccinimide (NHS) esters flanking an 11.4 Å spacer arm (Fig 1A). The NHS-esters react with primary amines of amino acids, typically lysines, but can also react with serines, tyrosines, and threonines ^22^ cross-linking amino acids within a 26-30Å proximity ^23^. DSS (0.25 mM) treatment of cells resulted in protein cross-linking, resulting in a characteristic molecular weight (MW) shift shown by SDS/PAGE and Western blotting (Fig 1B-C). Immunoprecipitation with anti-V5 antibodies resulted in the enrichment of PIP5Pase-V5 (Fig 1C). MS analysis from 11 experiments identified 158,718 total peptides, of which 41.4% were inter-links among different proteins, 0.7% were intra-links (mono or loop links), and 57.9% were not cross-linked (Fig 1D). A total of 50,593 cross-linked protein pairs were identified with a reproducibility index of 8 (±2.2), i.e., on average, a cross-link was reproduced in 8 experiments, and a mean of 13.2 (±14.1) cross-links per protein (Fig S1 and Table S1).

**Figure 1.**
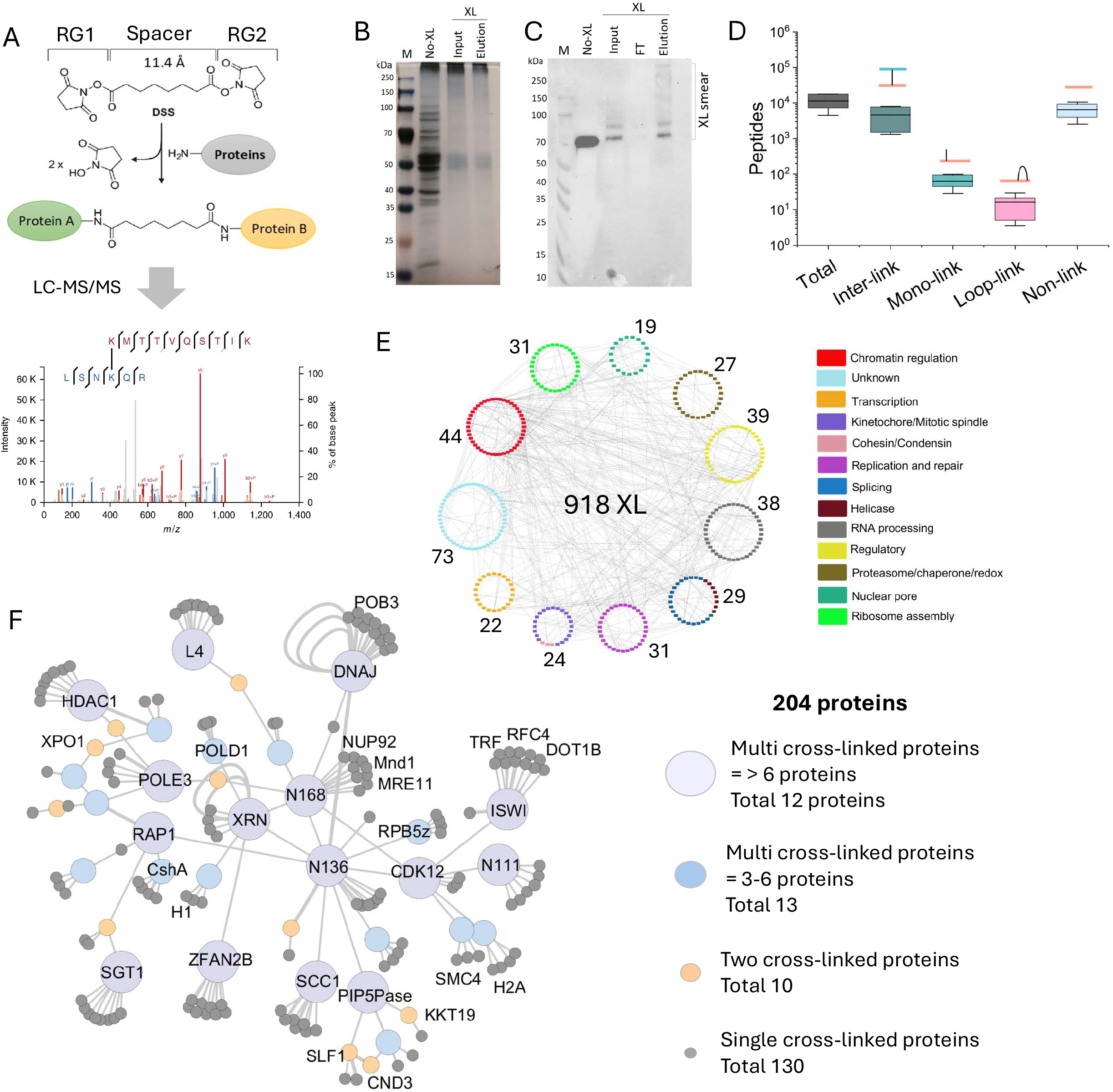
Chromatin regulatory protein network by XLMS. A) Diagram of DSS cross-linker, its reaction with proteins, and cross-linked amino acids (AAs) detection by LC-MS/MS. B-C) Silver-stained 10% SDS/PAGE (B) and Western blot (C) of immunoprecipitated V5-tagged PIP5Pase with α-V5 monoclonal antibodies (mAb) from a lysate of DSS cross-linked *T. brucei* bloodstream forms. Western blot probed with α-V5 mAbs. D) Total number of nuclear peptides detected by mass spectrometry and cross-link types. E) Cross-links of a subset (377) of nuclear proteins identified by PIP5Pase immunoprecipitation organized by functional categories. The number of proteins is indicated for each category. F-G) PIP5Pase cross-link interaction network. Edges represent cross-links, and nodes represent proteins. XL, cross-link; RG, NHS reactive group. Figures D, E and F show the combined data of 11 biological replicates.

The data revealed a nuclear cross-linked network of 494 proteins with 1,713 cross-links and 632 cross-link pairs involved in nucleic acid processes (Table S2), including chromatin regulation, kinetochore, transcription and splicing, nuclear pore, and regulatory proteins such as protein kinases and phosphatases (Fig 1E). Analysis of bonafide nuclear proteins revealed ~80% of them cross-linked with other nuclear proteins (Fig S1), indicating that most organellar cross-links occurred primarily before cell lysis. Analysis of PIP5Pase direct and counterpart cross-links revealed a subnetwork of 204 proteins with 750 cross-links (Fig 1F, Table S1 and S2). There were 25 proteins involved in multivalent cross-links, suggesting that some proteins may act as interaction hubs (Fig 1F). The network encompassed primarily proteins involved in DNA recombination, transcription, and chromatin-associated factors, including RAP1, HDAC1, ZCW1, ISW1, and DOT1B, likely reflecting PIP5Pase subnuclear locations and role in chromatin regulation ^18–20^.

### Mapping protein interacting domains with XLMS

Analysis of cross-linked peptides identified potential interacting domains among several proteins involved in chromatin-associated processes (Fig 2A-C). PIP5Pase cross-links were surface exposed and mainly distributed through its N-terminus (aa 1-150), which is consistent with the N-terminus containing a predicted disordered region and the C-terminus containing the 5-phosphatase catalytic domain (Fig 2B). It cross-linked with the N-terminus of KKT19, a kinetochore complex Cdc2-like protein kinase ^24^, and with the cohesin subunit protein SMC5-SMC6 complex localization factor protein 1 (SLF1), which functions in chromatin organization and DNA break and repair ^2,25,26^ (Fig 2A-B, Table S2). SLF1 cross-linked with condensin subunit 3 (CND3) and a putative BRCA2 and CDKN1A-interacting protein (BCIP1) (Fig 2B), which also functions in chromosome segregation. The data revealed cross-links in multiple chromatin-associated factors (Fig 2C), perhaps reflecting the spatial proximity of proteins involved in chromatin organization, transcription and splicing.

**Figure 2.**
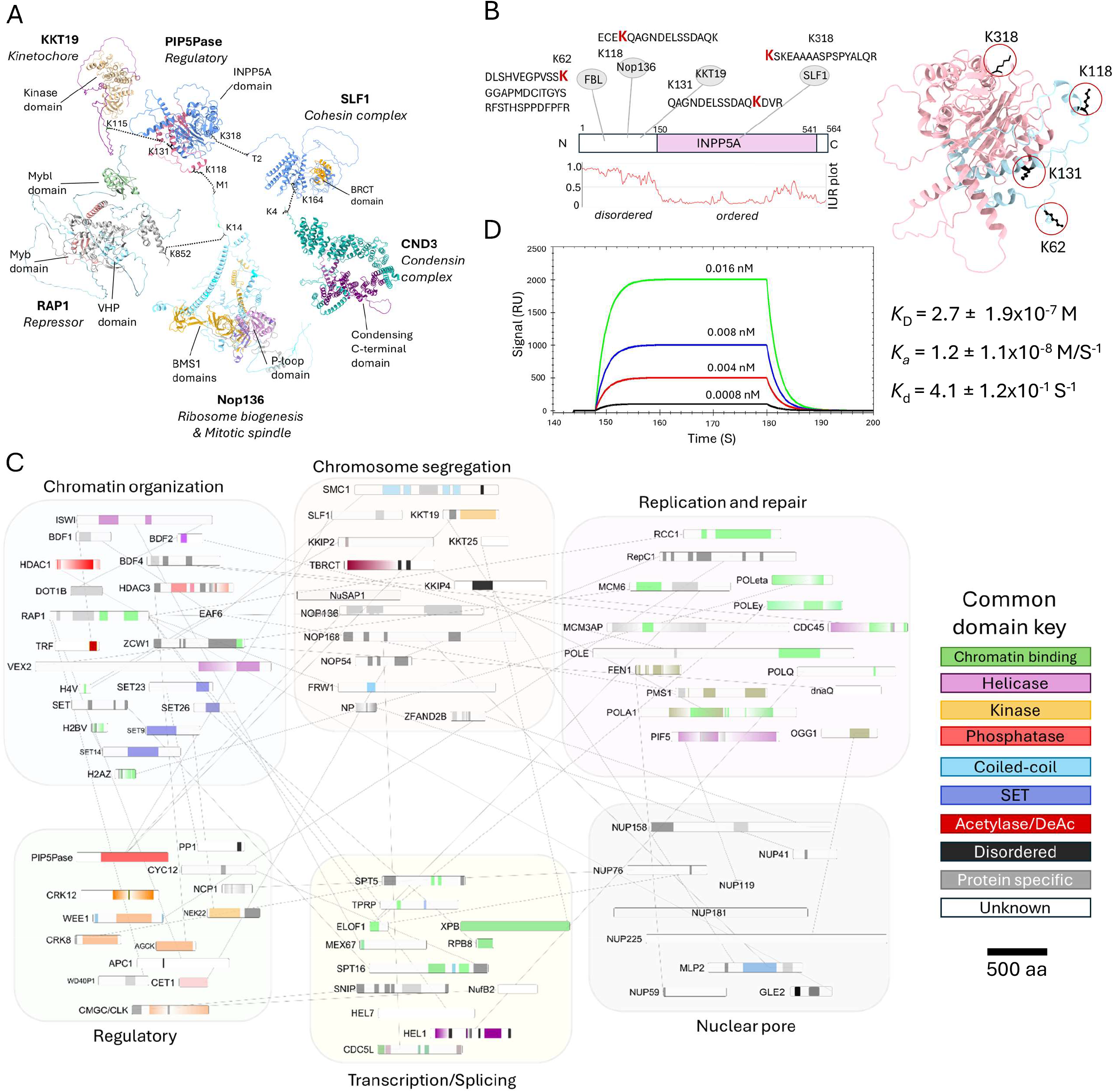
Mapping potential protein interacting domains with XLMS. A) Selected DSS cross-links between PIP5Pase (AlphaFold ID AF-Q385E2-F1-v4), KKT19 (AF-Q382T9-F1-v4), SLF1 (AF-Q57YW5-F1-v4), CND3 (AF-Q57ZK1-F1-v4), RAP1 (AF-Q387L4-F1-v4), and Nop136 (AF-Q384C0-F1-v4) indicated in the AlphaFold predicted protein structures. Dotted lines indicate cross-linked residues. Protein domains are indicated in colour. Protein function is indicated. B) PIP5Pase diagram and cross-linked residues. The AlphaFold predicted structure (AF-Q385E2-F1-v4) is shown on the right with cross-linked residues indicated. C) PIP5Pase cross-link network of selected chromatin-associated proteins showing cross-linked protein regions. See Table S2 for gene ID, product name abbreviations, and cross-linked residues. C) Native PIP5Pase-V5 and recombinant RAP1-His interactions by SPR. Data from three biological replicates. *K*_D_, equilibrium dissociation constant; *K*_d_, dissociation rate constant; *K*_a_, association rate constant. Data show mean ± standard deviation of the mean (SDM). IDR, intrinsically disordered regions. INPP5A, inositol polyphosphate-5-phosphatase A domain.

Our previous work suggested that PIP5Pase interacts with RAP1 ^18,20^. Analysis of RAP1 cross-links showed a distribution to its N- and C-terminus (aa 2-13 or aa 847-852) and distal from the villin headpiece (VHP) or the Myb and Myb-like domains (Fig 2C, Table S1), which we showed bind to PI(3,4,5)P3 and DNA, respectively ^19^. The protein Nop136, which has multiple disordered regions and functions in nucleolus assembly and chromosome segregation, cross-linked with RAP1 and PIP5Pase (Fig 2A-C), indicating spatial proximity among them. Binding kinetics by surface plasmon resonance (SPR) using *T. brucei* his-tagged RAP1 recombinantly produced in *E. coli* and V5-tagged PIP5Pase isolated from *T. brucei* ^18–20^ showed that PIP5Pase and RAP1 interacted directly with a *K*_D_ of 2.7 nM (± 1.9 nM), confirming their proximity by cross-link. The absence of a direct cross-link between the two proteins may result from the stringency of the *in vivo* cross-link approach, i.e., two cross-linkable amino acids (aa) within ~26-30Å proximity. Immunoprecipitation and mass spectrometry analysis of procyclic cells exclusively expressing a V5-tagged PIP5Pase allele ^19^ also identified RAP1 with 4.7 log2 enrichment (*p*-value < 0.001, Fig S2) and other proteins (Table S3), indicating conserved interactions in both parasite life stages. The XLMS data revealed a network of chromatin-binding proteins and their potential interacting domains, likely reflecting the dynamic interactions of chromatin-associated and regulatory processes *in vivo*.

### Interacting proteins at chromosome compartment boundaries

We postulated that the cross-links between chromatin-associated proteins reflect their spatial proximity due to chromatin’s three-dimensional organization. Thus, we performed Hi-C and ChIP-seq analysis in *T. brucei* bloodstream forms. *T. brucei* chromosomes are organized into core transcribed and subtelomeric repressed regions, with some chromosomes containing ESs (Fig 3A). We obtained ~155M chromatin contacts, ~118M intra- and ~37M inter-chromosomal contacts (Fig 3B, Table S4 and S5). We found that each chromosome contained about two or three compartments with an average length of 1.2 Mb (± 1 Mb, range of ~1-4 Mb), with core transcribed regions forming distinct compartments from silent subtelomeric regions (Fig 3C), analogous to A and B compartments, respectively ^1^. We found a higher number of intra-chromosomal contacts within subtelomeric compartments than in core compartments (Fig 3D), in agreement with previous findings ^7^. Moreover, we found that subtelomeric compartments of different chromosomes co-interacted with higher frequency than core compartments (Fig 3E), indicating that subtelomeric chromatin is spatially clustered, consistent with our observations by fluorescence *in situ* hybridizations ^20^. The compartments were further organized into multiple TADs, sub-TADs, and loops (Fig 3C, F-G), with the former averaging 571 Kb (± 479 Kb, *p*-value ≤ 0.05) and encompassing one to four PTUs (Fig 3G and 4G), whereas loops averaged 138 Kb (±182 Kb, *p-*value ≤ 0.05) (Fig 3F). The data indicate defined levels of intra and inter-chromosomal organization with silent and transcribed regions forming distinct compartments.

**Figure 3.**
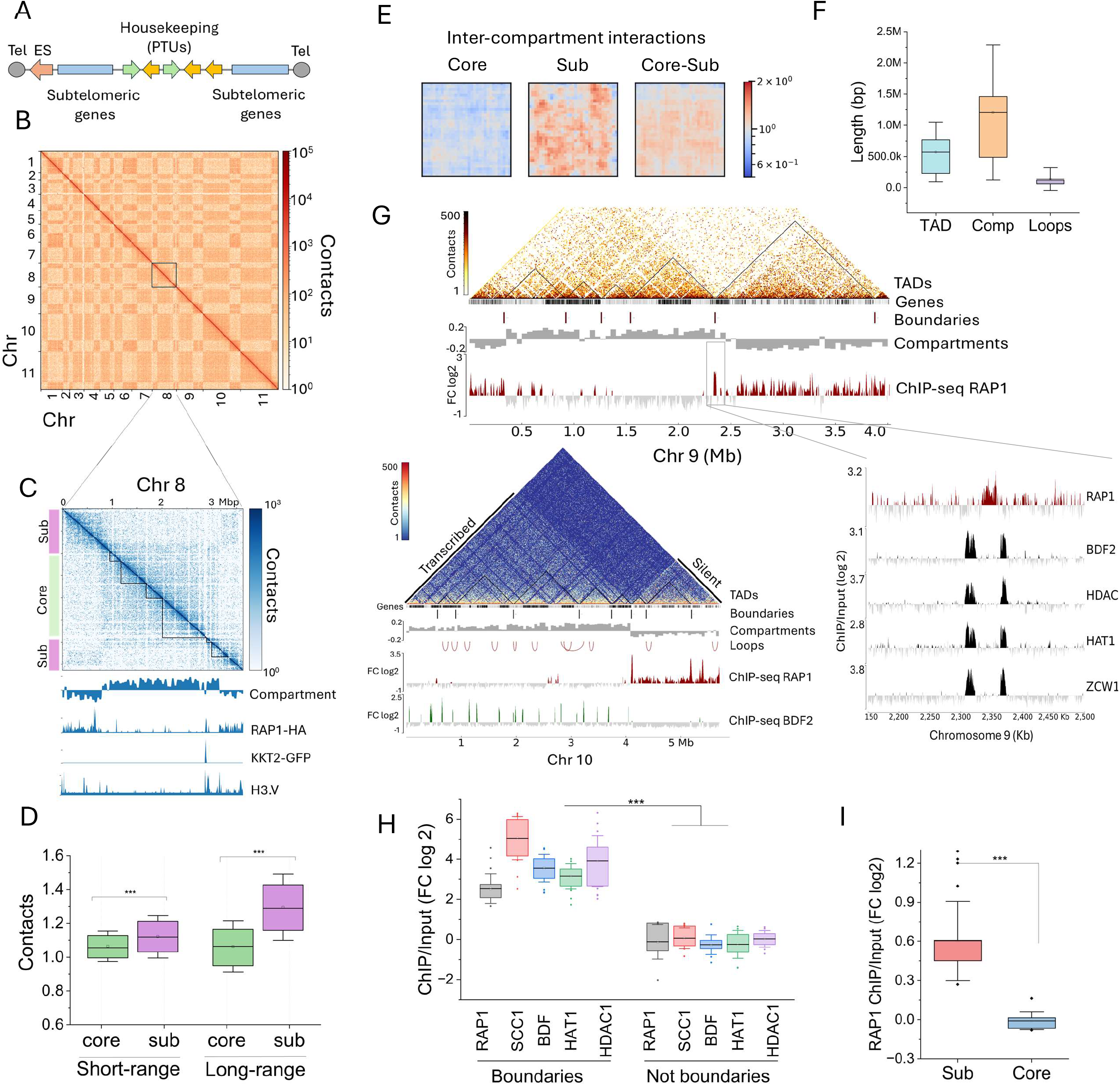
Chromosome spatial organization in *T. brucei* bloodstream forms. A) Diagram of *T. brucei* large chromosomes. PTUs, polycistronic units; Tel, telomere; ES, expression sites. B) Hi-C heatmap of intra- and inter-chromosomal contacts from chromosomes (Chr) 1 to 11. C) Hi-C heatmap of Chr 8 (top) with TADs (black lines) indicated. Compartments and ChIP-seq of RAP1-HA, KKT2, and H3.V are also indicated (bottom). A diagram of chromosome core and subtelomeres is shown on the left. D) Average short- and long-range intra-chromosomal interactions from all core and subtelomeric (sub) compartments. E) Average of all inter-chromosomal core and subtelomeric compartment interactions. F) Average length of TADs, compartments, and loops in bp. G) A Hi-C heatmap of Chr 9 (top) and 10 (bottom) interactions with compartments and TADs indicated. Compartments were assigned transcribed or silent based on RNA-seq data ^18,19^. ChIP-seq data of RAP1-HA, BDF2, HDAC, HAT1, and ZWC1 is shown at the Chr9 compartment boundary (~100 Kb segment). H) ChIP-seq enrichment of proteins in chromosome compartment boundaries. The average of all chromosome boundaries is shown. FC, fold-change of ChIP vs Input. Not boundaries signal was obtained from 10 Kb region flaking boundaries. I) ChIP-seq enrichment of RAP1-HA in subtelomeric vs core chromosome compartments. Data shows the average of all chromosomes. Core, core transcribed compartments; sub, subtelomeric silent compartments. D and E show interactions in median pixel values ^51^. Boxes in D, F, H, and I, show 25-75% data distribution; box line, the mean; and bars, the ± SDM. ^***^, *p-*value ≤ 0.001.

Our Hi-C and ChIP-seq revealed that RAP1 was markedly enriched at the boundaries of chromosome compartments separating subtelomeric from core regions (Fig 3G, H). Moreover, RAP1 spread over silent subtelomeric regions with a significant enrichment compared to core regions (Fig 3G, I). Notably, RAP1 chromatin binding correlates with silent subtelomeric compartments (Fig 3G), indicating that it might function to insulate and repress them. KKT2, enriched in centromeres ^7,24^, and H3.V were also found at the boundaries of some – but not all – compartments (Fig 3C), indicating that not all boundaries are centromeric (Fig 3C). Other chromatin-associated proteins, such as BDF2, HAT1, HDAC1, and ZCW1, which are at the limits of PTUs ^28^, were also enriched at compartment boundaries, typically flanking RAP1 binding sites, but were often depleted from subtelomeric regions (Fig 3G-H). The co-existence of these proteins at the compartment boundaries may explain their proximity as identified by XLMS, suggesting that they might contribute to insulating core from subtelomeric chromosome regions.

### Spatial chromatin compartment organization is essential for subtelomeric VSG gene silencing

The data indicate that the PIP5Pase network of chromatin-associated proteins may play a role in chromatin compartment organization. We used a conditional null PIP5Pase to determine if its knockdown affects chromatin spatial organization and gene expression within the compartments. We removed the endogenous PIP5Pase alleles and added a tetracycline (tet)-regulatable PIP5Pase, which can be turned on or off in the presence or absence of tet, respectively ^20^. We performed Hi-C in cells expressing PIP5Pase (tet +) or in which we knocked it down for 24h (tet −) (Fig 4A, Table S4 and S5). We found a remarkable decrease in short-range chromatin contacts and a slight increase in long-range contacts, including intra- and inter-chromosomal contacts (Fig 4B), the latter likely resulting from the disruption of chromosome organization. Analysis of chromosome compartments showed significant disruption of contacts within the compartments (Fig 4C-D). Moreover, inter-compartment interactions, i.e., contacts between compartments of different chromosomes, were also affected, primarily by disrupting subtelomeric compartment contacts (Fig 4E). The disruption of intra-chromosomal contacts in knockdown cells significantly affected the formation of TADs (Fig 4F), resulting in a decrease in TAD numbers or an increase in their size due to the loss of their boundaries (Fig 4G). Some chromosomes showed a decrease in the number of loops (Fig 4G), although loops appear to be less stable than TADs and compartments ^29–31^. The data indicates a role for PIP5Pase in regulating chromosome-level chromatin spatial organization.

**Figure 4.**
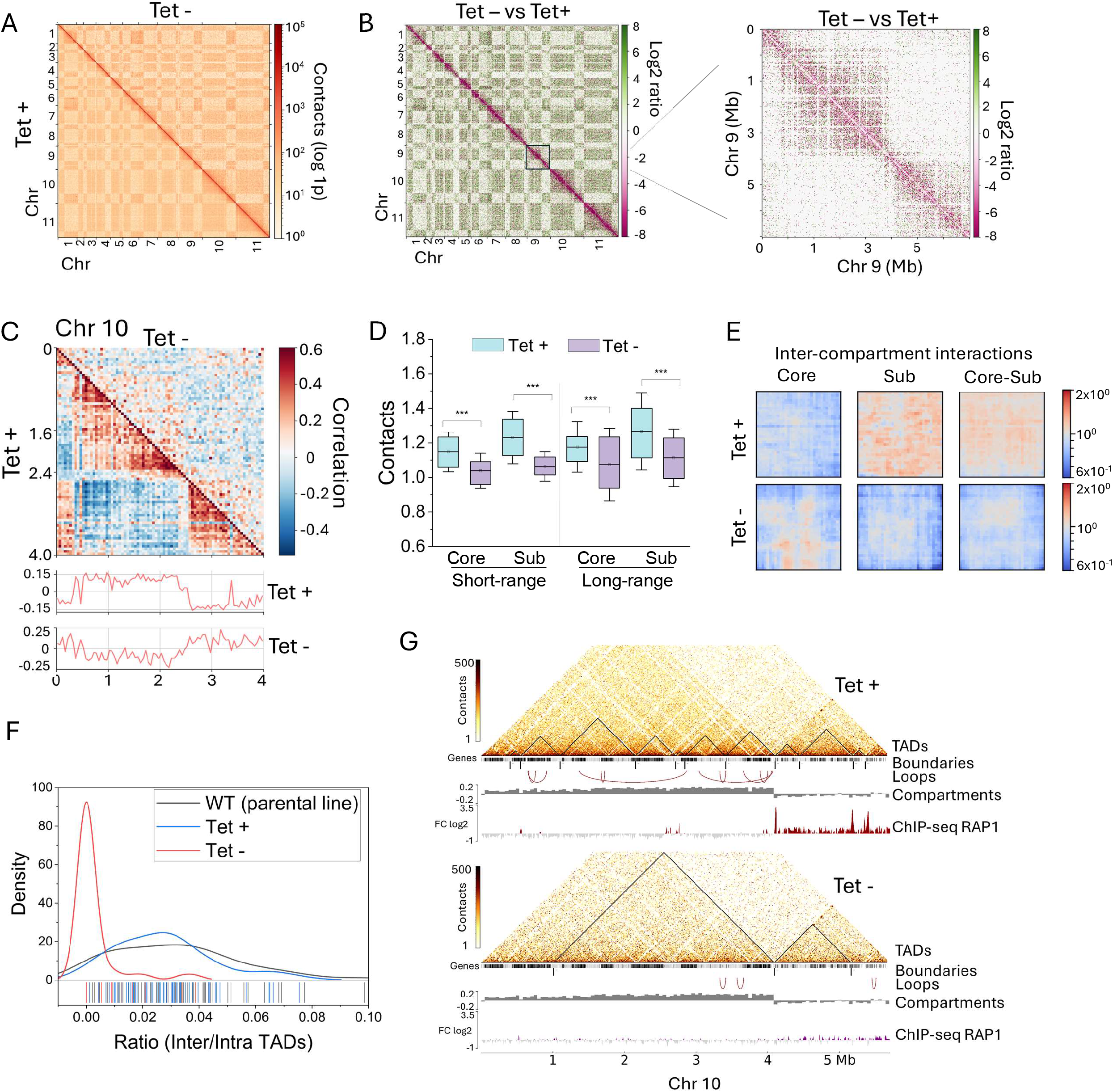
PIP5Pase regulates chromatin spatial organization. A) Hi-C heatmap of Chr 1-11 interactions *in T. brucei* expressing PIP5Pase (Tet +, lower left) or after 24h knockdown (Tet −, upper right). B) Heatmap of Chr 1-11 log2 ratio of interactions from PIP5Pase 24h knockdown (Tet −) versus PIP5Pase expressing cells (Tet +). On the right, Chr 9 log2 ratio of interactions from Tet – versus Tet + cells. C) Heatmap of Chr 10 interactions from cells expressing PIP5Pase (Tet +, lower left) vs 24h knockdown (Tet −, upper right). Compartment scores are shown below. D) Average short- and long-range intra-chromosomal contacts from all core and subtelomeric compartments in cells expressing PIP5Pase (Tet +) or after 24h knockdown (Tet −). E) Average of all inter-chromosomal core and subtelomeric compartment contacts in cells expressing PIP5Pase (Tet +) or after 24h knockdown (Tet −). F) Ratio of contacts between (Inter) and within (Intra) TADs in cells expressing PIP5Pase (Tet +) or after 24h knockdown (Tet −). Density, kernel density distribution of the data. X-axis bars show ratio data distribution. WT, single-marker 427 cells. G) Heatmap of Chr 10 chromatin contacts from cells expressing PIP5Pase (Tet +) or after 24h knockdown (Tet −). TADs, TAD boundaries, loops, compartments, and RAP1-HA ChP-seq are indicated. D and E show interactions in median pixel values ^51^. Boxes in D show 25-75% data distribution; box line, the mean; and bars, the ± SDM. ^***^, *p-*value ≤ 0.001.

To analyze if PIP5Pase controls RAP1 binding to subtelomeric compartments and thus their repression, we analyzed RAP1-HA ChIP-seq in cells expressing a catalytic inactive PIP5Pase, which fails to dephosphorylate nuclear PI(3,4,5)P3 ^18^ – an allosteric regulator of RAP1-DNA binding ^19^. RAP1 was abolished from chromatin compartment boundaries (Fig 4G, Fig 5A, B) and significantly decreased from subtelomeric silent compartments in cells expressing catalytic inactive PIP5Pase (Fig 4G, Fig 5A and C). Immunofluorescence analysis of HA-tagged RAP1 showed a broader nuclear distribution of RAP1 occupying a 6-fold area increase in cells expressing mutant compared to wildtype PIP5Pase (Fig 5D, E), perhaps reflecting loss of RAP1-DNA association. Moreover, RNA-seq comparing PIP5Pase catalytic mutant versus wildtype (WT) showed that subtelomeric compartment genes were significantly de-repressed, resulting in the expression of subtelomeric VSGs (Fig 5A, F). The data indicates that PIP5Pase controls chromatin compartment organization and transcriptional repression of subtelomeric VSG genes via RAP1 association with compartment boundaries and subtelomeric regions.

**Figure 5.**
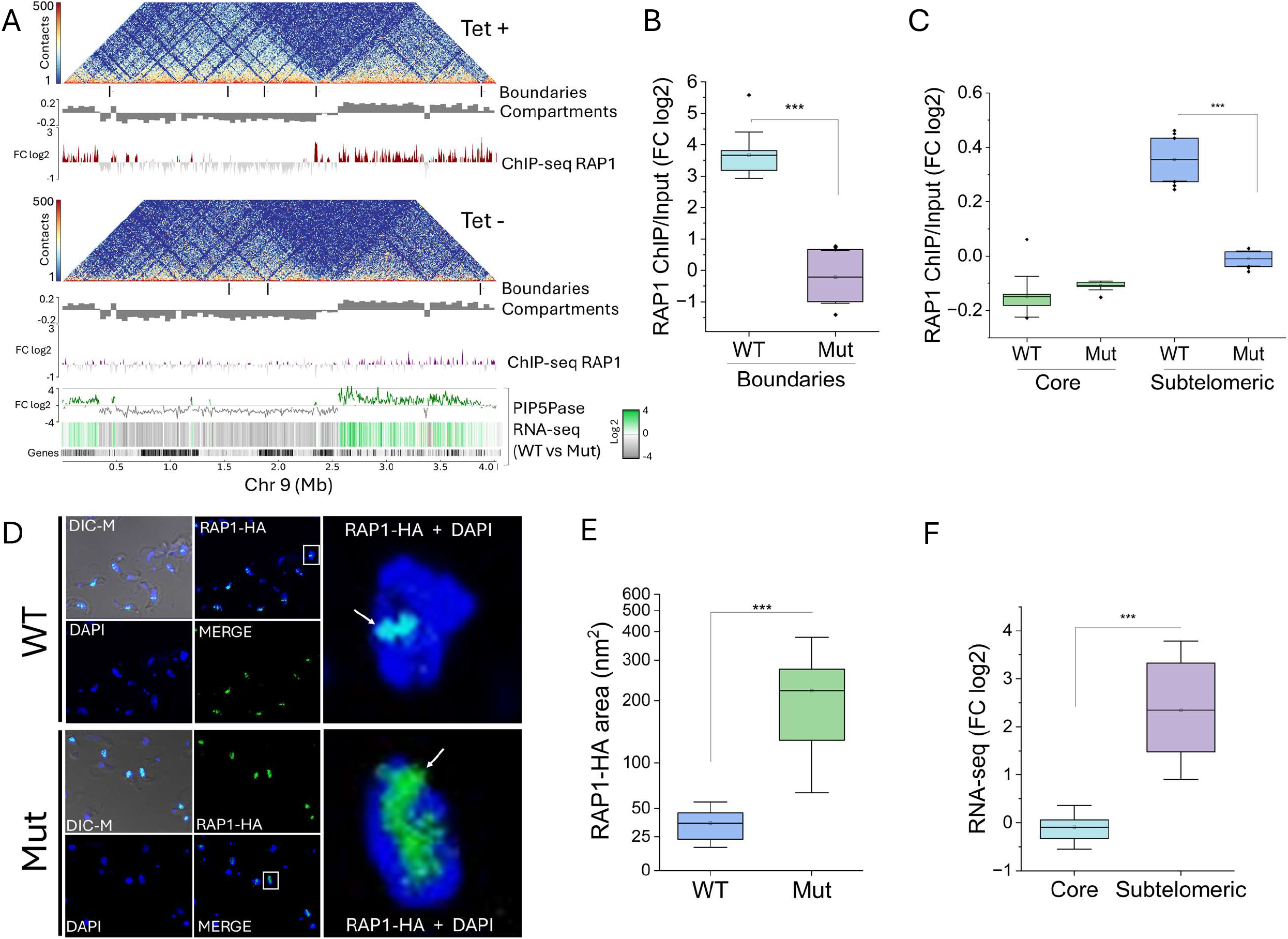
PIP5Pase and RAP1 represses VSG-rich subtelomeric compartments. A) Heatmap of Chr 9 chromatin contacts in cells expressing PIP5Pase (Tet +) or after its 24h knockdown (Tet −). Below the heatmap are TAD boundaries, compartments, and RAP1-HA ChIP-seq in cells exclusively expressing PIP5Pase (WT) or its catalytic inactive mutant (Mut, D362A/N360A), and RNA-seq heatmap from cells exclusively expressing PIP5Pase (WT) versus its catalytic inactive mutant (Mut). B-C) RAP1-HA ChIP-seq enrichment in compartment boundaries (B) and subtelomeric and core compartments (C) in cells exclusively expressing WT or Mut PIP5Pase. FC, fold-change of RAP1-HA ChIP-seq vs Input. Data shows the average of all chromosome boundaries and compartments. D) Confocal microscopy of HA-tagged RAP1 (green) in *T. brucei* exclusively expressing WT or Mut PIP5Pase for 24h. DNA stained with DAPI (blue). E) Quantification of the nuclear area occupied by RAP1-HA from Fig D. N = 458 nuclei. F) Change in the expression of genes from subtelomeric and core compartments in cells exclusively expressing WT or Mut PIP5Pase for 24h. Data shows the average of all genes within all core or all subtelomeric chromosome compartments. Boxes in B, C, E and F show 25-75% data distribution; box line, the mean; bars, ± SDM; and dots are outliers. ^***^, *p-*value ≤ 0.001.

## Discussion

We found that the spatial organization of chromatin into chromosome compartments is essential for silencing subtelomeric genes in *T. brucei*. In this organism, chromosomes are organized into two distinct compartments: transcribed core with housekeeping genes and silent subtelomeric with primarily VSG genes. Using XLMS, we uncovered chromatin-associating factors contributing to this chromosome spatial organization. Among these factors, we show that RAP1 marks the compartment boundaries and functions in silencing subtelomeric genes. Subtelomeric compartment silencing also depends on PIP5Pase activity, which controls chromosome contacts and RAP1 binding to subtelomeric DNA. The data points to a regulatory mechanism that spatially segregates transcribed and repressed chromatin and represses hundreds of VSG genes. Furthermore, it shows a role for phosphoinositides in the control of genome spatial organization.

In addition to compartments, we show that *T. brucei* chromosomes are further sub-organized into TADs, sub-TADs, and loops with length ranges correlating with other eukaryotes ^1–3^. The chromosome spatial organization likely reflects the cell’s need to control transcription, which in some organisms occurs by placing co-regulated genes, promoters and enhancers in proximity ^2,6^. *T. brucei* lacks RNA polymerase II transcriptional control, and genes within PTUs of core chromosome regions are co-transcribed ^32^. On the other hand, the silent VSG genes are physically separated from core genes and arranged at the subtelomeres. The spatial compartmentalization of core and subtelomeric regions might function to isolate repressed from transcribed regions. We also show that multiple subtelomeric compartments co-interact, further supporting a genome-wide spatial organization that separates transcribed from repressed chromatin. *T. brucei* genome might have evolved physical and spatial segregation of silent and active genes to compensate for the lack of transcriptional control.

Analysis of XLMS showed that the PIP5Pase interaction network comprises proteins involved in chromatin organization, transcription, splicing, DNA repair and recombination, among others. The cross-linking of these proteins within a short ~26-30 Å range might have resulted from their direct binding and/or the spatial proximity of their chromatin binding sites. For example, factors involved in transcription, DNA replication, or chromatin organization may co-exist associated with the same chromosome regions but play distinct roles. Notably, the cross-link indicates the proximity of proteins and not their direct interaction. Conversely, the lack of cross-link does not imply a lack of interaction due to the approach’s stringency requiring two cross-linkable amino acids within 26-30Å^23^, as we demonstrated for PIP5Pase and RAP1. The presence of RAP1 and other chromatin-associated factors, e.g., BDF2, HDAC1, HAT1, and ZWC1, at the compartment boundaries suggest a role for these proteins in chromosome compartment organization. The boundaries and proteins associated thereof might function to delimit transcribed and repressed states of the chromatin. The removal of RAP1 from boundaries and subtelomeric compartments after PIP5Pase inactivation concurred with disruption in chromatin contacts and subtelomeric genes transcriptional activation, indicating that spatial compartment organization and both proteins are required to silence subtelomeric chromatin.

Chromosome compartments are likely formed every cell cycle after DNA replication. PIP5Pase might function to control RAP1 binding to compartment boundaries and subtelomeric regions during compartment assembly through/after DNA replication, and the failure of RAP1’s association with these regions may affect compartment assembly and their repression. PIP5Pase knockdown also disrupted the TADs, which could be a consequence of the misassembled compartments or a direct role of this enzyme in controlling the formation of TADs, perhaps via its association with the SLF1 subunit of the cohesin complex. We showed that disruption of PIP5Pase activity also results in VSG switching by transcriptional switching among ESs and by recombination of subtelomeric VSG genes within ESs ^19,33^. We found here that subtelomeric compartments of multiple chromosomes co-interact, which may help isolate the silent subtelomeric genes. Disruption of this chromosome organization by PIP5Pase knockdown increases unspecific contact among various chromosomes, including with telomeric ESs, which may favour VSG gene recombination. PIP5Pase regulation of these processes implies that they occur in regions containing phosphoinositides. Phosphoinositides are synthesized in the endoplasmic reticulum and Golgi and redistributed to other cellular compartments, including the nucleus, where they are modified ^20,34^. PI(3,4,5)P3 association with RAP1 might result from this protein’s proximity to nuclear membranes through interactions with nuclear lamina proteins ^18^, such as NUP-1, which knockdown also affects subtelomeric gene silencing ^35^. Alternatively, these metabolites may associate with the chromatin via nucleoplasmic lipid droplets containing phosphoinositides ^36^. In any case, our data indicate a lipid signalling control of chromatin spatial organization. ^36^.

Our results suggest that *T. brucei* chromosomes evolved to physically and spatially segregate transcriptionally active from repressed genes via the assembly of chromosome compartments. Moreover, it shows that RAP1, along with other chromatin factors such as BDF2, HDAC, HAT1, etc., marks the compartment boundaries, whereas RAP1 also spreads over subtelomeric regions and represses subtelomeric VSG genes. Given that RAP1 also associates with silent telomeric ESs ^19^, it may function as a general transcriptional repressor in this organism. The regulation of chromosome topology by PIP5Pase indicates an epigenetic role for phosphoinositides in controlling genome spatial organization, a process that may be conserved in other eukaryotes.

### Materials and Methods

### Cell culture and cell lines

*T. brucei* 427 derived cell lines were maintained in HMI-9 medium supplemented with 10% (vol/vol) fetal bovine serum (FBS) at 37°C with 5% CO_2_. *T. brucei* expressing the C-terminally V5-tagged PIP5Pase or HA-tagged RAP1 were generated by transfection of pMOTag2H or pMOTag3V5 into one of their respective alleles, as previously described ^37^, and maintained in the presence of 0.1 μg/mL of puromycin. Conditional null PIP5Pase in bloodstream forms ^20^, or modified lines to exclusively express wildtype or mutant (D360A/E362A) PIP5Pase ^18^ were maintained at 10 μg/mL of zeocin and 0.1 μg/mL of nourseothricin N-acetyl transferase. Conditional null PIP5Pase in procyclics ^19^ were maintained at 27°C in SDM-79 (Wisent) medium containing 10% FBS and penicillin/streptomycin (Thermo Fisher Scientific), supplemented with 15 μg/mL of G418, 25 μg/mL of hygromycin and 2.5 μg/mL of zeocin. Tetracycline at 0.5 μg/mL was added to CN PIP5Pase cultures to induce PIP5Pase expression.

### *In vivo* chemical cross-linking of *T. brucei* bloodstream forms

*T. brucei* cells were grown in five litres of HMI-9 medium as described above. The cells were washed five times in phosphate-buffered saline (100 mM sodium phosphate dibasic anhydrous, 100 mM sodium phosphate monobasic anhydrous, 145 mM sodium chloride) supplemented with 6 mM glucose (PBS-G). Cells were centrifuged at 4,000 rpm then resuspended in 500ml of PBS-G. DSS (ProteoChem) dissolved in dimethylsulfoxide (DMSO) (Sigma) was added at 0.1 mM to the cell suspension. The mix was incubated for 10 minutes with gentle shaking at 37°C. The reaction was quenched by adding 50 mM glycine pH 9 and incubated for 15 minutes at room temperature (RT). Cells were spun down at 4,000 rpm, the pellet resuspended in 10ml of lysis buffer (50 mM Tris, 150 mM NaCl, 1% Triton X-100, 0.5% sodium deoxycholate, 0.1% NP-40, 2x protease inhibitor cocktail (Roche), pH 8.0), and then incubated for 30 minutes gently rotating at 4°C. Lysate was centrifuged at 14,000 rpm for 20 minutes at 4°C, and soluble proteins were collected from the supernatant. Proteins were analyzed in 10% SDS-PAGE and by Western blot for PIP5Pase-V5 expression before immunoprecipitations.

### Immunoprecipitation of PIP5Pase from cross-linked cells

Monoclonal antibodies (mAbs) α-V5 (ABclonal Inc.) were cross-linked to protein G magnetic beads (Cytiva) as previously described ^18^. Fifty microliters of Protein-G-α-V5 mAbs (20 μg of α-V5) were incubated with 10 mL of protein lysate and incubated overnight, gently rotating at 4°C. Afterward, the mixture was washed in wash buffer (50 mM Tris-HCl, 300 mM NaCl, 0.1% NP40, 2x protease inhibitor cocktail (Roche), pH 8.0) using a magnetic stand. The proteins were eluted twice (200 μl each) in 6 M urea/100 mM glycine pH 2.9. Ten percent of the elute was collected for SDS-PAGE and Western analysis, and the remaining was kept at −80°C for mass spectrometry analysis. For procyclic cells expressing V5-tagged PIP5Pase, 2.0×10^10^ cells were harvested and lysed ^18^, and immunoprecipitated with α-V5 monoclonal antibodies as indicated above.

### Protein digestion and mass spectrometry

Protein digestion was adapted from ^38^. Briefly, six volumes of cold acetone (−20°C) were added to immunoprecipitation eluate, vortexed and incubated for 1 hour at −20°C for protein precipitation. Afterward, protein samples were centrifuged at 14,000 x g for 30 minutes at 4°C. The acetone was decanted and air-dried for 15 minutes, and pellets were resuspended in 10μl of 6M urea. Then, 5 μl of reducing solution (10 mM DTT in 100 mM NH_4_CO_3_) were added, and samples were vortexed and incubated for 1 hour at room temperature. Samples were spun down (3,000 x g for 15 seconds), 3 μl of alkylation solution (50 mM iodoacetamide in 100 mM NH_4_CO_3_) were added, and samples were incubated in the dark for 30 minutes at RT. Next, 3 μl of reducing solution was added to neutralize the reaction. Lys-C/trypsin mix (Promega) was added to the protein in a ratio of 1:50 (protein: protease amounts in μg), mixed and incubated for 4 hours at 37°C. The sample was diluted six-fold with 50 mM Tris-HCL (pH 8.0) to bring the concentration of UREA to 1 M and the reaction incubated overnight at 37°C. The reaction was terminated by adding 13.06 M trifluoroacetic acid to a final concentration of 1%. Any particulate material was removed by centrifugation at 14,000 × g for 10 minutes. The supernatant was analyzed in 10% SDS/PAGE to verify digestion. The samples were analyzed in Orbitrap LC-MS at the McGill University Proteomics Centre. Briefly, peptides were solubilized in 0.1% aqueous formic acid, loaded onto a Thermo Acclaim Pepmap (Thermo, 75μM ID X 2cm C18 3μM beads) precolumn and then onto an Acclaim Pepmap Easyspray (Thermo, 75μM X 15cm with 2μM C18 beads) analytical column separation using a Dionex Ultimate 3000 uHPLC at 230 nl/min with a gradient of 2-35% organic (0.1% formic acid in acetonitrile) over 2 hours. Peptide analysis was done using a Thermo Orbitrap Fusion mass spectrometer operating at 120,000 resolution (FWHM in MS1) with HCD sequencing (15,000 resolution) at top speed for all peptides with a charge of 2+ or greater. The raw data was converted to mzML (MSConvert) for downstream processing and analysis.

### Cross-link mass spectrometry data analysis

Peptide and protein identification was done using the Trans Proteomic Pipeline (http://www.tppms.org/) ^39^. Thermo RAW files were first converted to mzML with MSConvert, and cross-linked peptides were identified using Kojak version 2.0.3 ^40^ using default parameters except for top_count adjusted to 50, min_peptide_mass to 600, max_peptide_mass to 8000, ppm_tolerance_pre to 25, MS1_resolution and MS2_resolution adjusted to 60000 and 50000, respectively, min_peptide_score to 0.5, min_spectrum_peaks to 12 and max_spectrum_peaks to 0. The search was done against *T. brucei* 927 strain reference genome v9.0 predicted protein sequence database. A false discovery rate (FDR) threshold of 0.05 with precursor mass tolerance of 10 ppm and two missed cleavages allowed were used. Cross-linking data was validated using Percolator ^40^. Protein interactions were visualized using Cytoscape ^41^, ProxL ^42^ and xiNET ^43^. For procyclic cells, data was processed using MaxQuant with default parameters and Perseus for statistical analysis^44^.

### Western Blotting

Western blot analysis was performed as previously described with modifications ^18^. Briefly, lysates of *T. brucei* were prepared in lysis buffer containing 50 mM Tris-HCl pH 8.0, 300 mM NaCl, 0.1% NP40, 1% Triton-X-100, 2x protease inhibitor cocktail (Roche) and 0.5% sodium deoxycholate. Cleared lysate was mixed in 4x laemmli loading buffer containing with 2.85 M β-mercaptoethanol, and heated for 5 minutes at 95°C. Proteins were resolved in 10% gels and transferred to a polyvinylidene difluoride (PVDF) membrane. The membrane was probed for 2 hours at RT (or overnight at 4°C) with mAbs α-HA (ABclonal) or mAb α-V5 (ABclonal) diluted 1:5,000 in 10% skimmed milk (Bio-Rad) dissolved in PBS 0.05% Tween 20. The membranes were incubated with goat anti-mouse IgG (H+L)-HRP 1:2000 (Bio-Rad) and developed by chemiluminescence using a GelDoc Imaging System (Bio-Rad).

### Immunofluorescence assay

*T. brucei* were grown to mid-log phase, washed three times in PBS-G by centrifugation at 4,000 rpm for 5 min, fixed with 2% paraformaldehyde (Electron Microscopy Sciences, PA) in PBS and adhered to poly-l-lysine-treated 2-mm cover slips (Fisher). Cells were permeabilized with 0.2% NP-40 (Sigma Aldrich) in PBS for 10 min and blocked for 1□h at RT in blocking buffer (10% nonfat dry milk diluted in PBS). Cover slips were incubated for 2□h at RT in either mAb α-HA 3F10 (Invitrogen), or mouse mAb α-V5 (Thermo Life Technologies) each at a ratio of 1:500 diluted in blocking buffer, followed by goat α-mouse IgG (H+L)–Alexa Fluor 488 (Thermo Fisher Scientific) in 10% nonfat dry milk diluted in PBS. Slides were mounted with Fluoromount G mounting medium with 4’,6-diamidino-2-phenylindole (DAPI) (Biotium). Images were acquired with Zeiss Confocal Laser Microscope and analyzed with Zen Microscopy software (Zeiss).

### Hi-C and computational analysis

Hi-C was performed in *T. brucei* bloodstream forms single marker 427 or conditional null PIP5Pase. PIP5Pase was knocked down for 24h (tet −) and compared to cells expressing PIP5Pase (tet +). 3×10^8^ cells were fixed in 1% paraformaldehyde in PBS for 10 minutes at RT and quenched with 0.2M glycine for 5 minutes. Cells were washed in PBS by centrifugation at 4,000xg for 10 minutes. Pellets were processed for Hi-C adapted from ^45^. Briefly, cells were lysed, and nuclear pellets were extracted and digested in NEB Buffer 2 with 100 units of *Mbo* I (New England Biolabs) at 37°C overnight with rotation. After *Mbo* I heat inactivation at 62°C for 20 minutes, DNA ends were filled with 1 mM biotinylated dATP (Thermo Fisher Scientific) and 10 mM dCTP, dGTP, dTTP with 50 units DNA polymerase I Klenow fragment (New England Biolabs) for 4h at 23°C. DNA ends were ligated with 4,000 units of T4 DNA ligase (New England Biolabs) at 16°C overnight, shaking at 80 rpm. Lysate was digested with 1.1 mg of proteinase K (New England Biolabs) at 63°C for 4h, and DNA was extracted by phenol:chloroform:isoamyl alcohol (25:24:1, v/v) method. The DNA was sonicated using a Covaris M220 ultrasonicator (50 peak incidence power, 20% duty factor, 200 cycles per burst, time 180 seconds). DNA fragments ranging from 100-450 bp were size selected from 1% agarose gel/TBE, end-repaired and A-tailed with Ultra II End Prep kit (New England Biolabs) and DNA-sequencing library prepared with NEBNext oligos for Illumina (New England Biolabs). Libraries were sequenced at GenomeQuebec. Raw fastq files were mapped to the *T. brucei* 427 genome ^7^ using bwa-mem ^46^, and DNA contacts were obtained using pairtools ^47^. Pair files were converted to cool files using cooler ^48^, and the matrixes of three biological replicates were combined. Compartments were analyzed with Fanc ^49^ and TADs and loops with HiCExplorer ^50^. Comparison between tet + vs tet – was performed using HiCExplorer. Compartment analysis and quantification were performed with Pentad ^51^. ChIP-seq and RNA-seq data were obtained from ^19,24,28^, and analyzed as previously described ^19^. All scripts are available at https://github.com/cestari-lab/lab_scripts.

### Surface plasmon resonance

Ten μg/ml of RAP1-His (110 nM) were diluted in binding buffer (20 mM HEPES [(4-(2-hydroxyethyl)-1-piperazineethanesulfonic acid)], 150 mM NaCl, 10 mM KCl, 10 mM MgCl2, 10 mM CaCl2 and 0.2% NP-40) for binding assays. The NTA-sensor (Nicoya) surface was cleaned in a solution of 10 mM HCl and 350 mM EDTA solution, then activated in 40 mM NiCl_2_ solution. RAP1-His was immobilized in the NTA sensor in binding buffer, followed by sensor blocking in binding buffer supplemented with 0.5% BSA. PIP5Pase-V5 diluted in binding buffer was added in various concentrations (0.0008 nM, 0.004, 0.008, 0.016 nM) for binding kinetic analysis. The sensor surface was regenerated for every binding assay using 500 mM Imidazole diluted in Nanopure MilliQ water. Reactions were prepared in 200 μL volume in 96-well plate and run in the OpenSPR-XT (Nicoya). Data was recorded and analyzed using the Traceviewer software (Nicoya).

### Resource availability

RNA-seq and ChIP-seq sequencing data are available in the Sequence Read Archive (SRA) with the BioProject identification PRJNA934938. Hi-C data is available in the SRA with BioProject identification PRJNA1198910. The mass spectrometry proteomics data have been deposited to the ProteomeXchange Consortium via the PRIDE partner repository with the dataset identifiers PXD059635 and PXD059549. Codes used for data analysis are available at https://github.com/cestari-lab/lab_scripts.

## Acknowledgements

This research was partly enabled by computational resources provided by Calcul Quebec (https://www.calculquebec.ca/en/) and the Digital Research Alliance of Canada (alliancecan.ca). We thank Dr. Suzanne McDermott (Center for Global Infectious Disease Research, Seattle Children’s) for reading the manuscript and providing valuable suggestions.

## Author’s contributions

T.I. performed cross-link and mass spectrometry, SPR protein binding assays, and microscopy experiments; L.B.A. optimized Hi-C protocol, performed Hi-C and Pore-C, designed Hi-C and Pore-C computational pipeline, analyzed Pore-C dataset, and performed ChIP-seq analysis; O.K. optimized pore-C protocol and preliminary data analysis; I.C. designed research, generated cell lines, performed Hi-C and ChIP-seq computational analysis. I.C. obtained research funds, supervised the research, and wrote the manuscript. All authors read and revised the manuscript.

## Supplemental figures

**Figure S1. Analysis of XL-MS dataset**. A) Graph shows the reproducibility of the XL-MS dataset and indicates a reproducibility index (mean) 7.9 (± 2.2), i.e., on average, a protein was detected at least on eight different biological replicates. B) Distribution of cross-links per protein. It shows a mean of 13.23 (± 14.1) cross-links per protein. C) Top, diagram for the analysis of cross-links that likely occurred *in vivo* or after cell lysis. NE, nuclear envelope. Bottom, quantification of cross-links of bonafide nuclear proteins with other nuclear (Nuc-Nuc), cytoplasmic (Nuc-Cyt), mitochondrial (Nuc-Mit), endoplasmic reticulum/Golgi apparatus (Nuc-Golg).

**Figure S2.** Identification of PIP5Pase interacting proteins in *T. brucei* procyclic forms. A) 10% SDS/PAGE Coomassie stained shows lysate and V5-tagged PIP5Pase immunoprecipitated with anti-V5 antibodies. FT, flow through. B) Western blotting of samples shown in A with anti-V5 antibodies. Asterisks indicate PIP5Pase migration (~62 KDa). C) Enrichment analysis of PIP5Pase-V5 interacting proteins. In grey, non-significant interactions; in purple, significant interactions with p-value <0.05 and fold-change (FC) > 2. D) Diagram of selected PIP5Pase interacting proteins (solid line). The dotted lines indicate proteins identified by XL-MS. See Table S3 for a list of interacting proteins.

## Supplemental Information List

**Figure S1 and Figure S2**.

**Table S1**. Excel file containing proteins identified by cross-linking and mass spectrometry.

**Table S2**. Excel file containing proteins in the nuclear PIP5Pase interacting network identified by cross-linking and mass spectrometry.

**Table S3-S5**.

## References

1. Lieberman-Aiden, E., van Berkum, N.L., Williams, L., Imakaev, M., Ragoczy, T., Telling, A., Amit, I., Lajoie, B.R., Sabo, P.J., Dorschner, M.O., et al. (2009). Comprehensive mapping of long-range interactions reveals folding principles of the human genome. Science 326, 289–293. 10.1126/science.1181369.

2. Rowley, M.J., and Corces, V.G. (2018). Organizational principles of 3D genome architecture. Nat Rev Genet 19, 789–800. 10.1038/s41576-018-0060-8.

3. Dixon, J.R., Selvaraj, S., Yue, F., Kim, A., Li, Y., Shen, Y., Hu, M., Liu, J.S., and Ren, B. (2012). Topological domains in mammalian genomes identified by analysis of chromatin interactions. Nature 485, 376–380. 10.1038/nature11082.

4. Cai, Y., Zhang, Y., Loh, Y.P., Tng, J.Q., Lim, M.C., Cao, Z., Raju, A., Lieberman Aiden, E., Li, S., Manikandan, L., et al. (2021). H3K27me3-rich genomic regions can function as silencers to repress gene expression via chromatin interactions. Nat Commun 12, 719. 10.1038/s41467-021-20940-y.

5. Nora, E.P., Lajoie, B.R., Schulz, E.G., Giorgetti, L., Okamoto, I., Servant, N., Piolot, T., van Berkum, N.L., Meisig, J., Sedat, J., et al. (2012). Spatial partitioning of the regulatory landscape of the X-inactivation centre. Nature 485, 381–385. 10.1038/nature11049.

6. Rodriguez-Carballo, E., Lopez-Delisle, L., Willemin, A., Beccari, L., Gitto, S., Mascrez, B., and Duboule, D. (2020). Chromatin topology and the timing of enhancer function at the HoxD locus. Proc Natl Acad Sci U S A 117, 31231–31241. 10.1073/pnas.2015083117.

7. Muller, L.S.M., Cosentino, R.O., Forstner, K.U., Guizetti, J., Wedel, C., Kaplan, N., Janzen, C.J., Arampatzi, P., Vogel, J., Steinbiss, S., et al. (2018). Genome organization and DNA accessibility control antigenic variation in trypanosomes. Nature 563, 121–125. 10.1038/s41586-018-0619-8.

8. Cestari, I., and Stuart, K. (2018). Transcriptional Regulation of Telomeric Expression Sites and Antigenic Variation in Trypanosomes. Curr Genomics 19, 119–132. 10.2174/1389202918666170911161831.

9. Cross, G.A. (1975). Identification, purification and properties of clone-specific glycoprotein antigens constituting the surface coat of Trypanosoma brucei. Parasitology 71, 393–417. 10.1017/s003118200004717x.

10. Vickerman, K., and Luckins, A.G. (1969). Localization of variable antigens in the surface coat of Trypanosoma brucei using ferritin conjugated antibody. Nature 224, 1125–1126. 10.1038/2241125a0.

11. Ferguson, M.A., Homans, S.W., Dwek, R.A., and Rademacher, T.W. (1988). Glycosyl-phosphatidylinositol moiety that anchors Trypanosoma brucei variant surface glycoprotein to the membrane. Science 239, 753–759. 10.1126/science.3340856.

12. Dakovic, S., Zeelen, J.P., Gkeka, A., Chandra, M., van Straaten, M., Foti, K., Zhong, J., Vlachou, E.P., Aresta-Branco, F., Verdi, J.P., et al. (2023). A structural classification of the variant surface glycoproteins of the African trypanosome. PLoS Negl Trop Dis 17, e0011621. 10.1371/journal.pntd.0011621.

13. Navarro, M., and Gull, K. (2001). A pol I transcriptional body associated with VSG mono-allelic expression in Trypanosoma brucei. Nature 414, 759–763. 10.1038/414759a.

14. Faria, J., Glover, L., Hutchinson, S., Boehm, C., Field, M.C., and Horn, D. (2019). Monoallelic expression and epigenetic inheritance sustained by a Trypanosoma brucei variant surface glycoprotein exclusion complex. Nat Commun 10, 3023. 10.1038/s41467-019-10823-8.

15. Glover, L., Hutchinson, S., Alsford, S., and Horn, D. (2016). VEX1 controls the allelic exclusion required for antigenic variation in trypanosomes. Proc Natl Acad Sci U S A 113, 7225–7230. 10.1073/pnas.1600344113.

16. Lopez-Escobar, L., Hanisch, B., Halliday, C., Ishii, M., Akiyoshi, B., Dean, S., Sunter, J.D., Wheeler, R.J., and Gull, K. (2022). Stage-specific transcription activator ESB1 regulates monoallelic antigen expression in Trypanosoma brucei. Nat Microbiol 7, 1280–1290. 10.1038/s41564-022-01175-z.

17. Yang, X., Figueiredo, L.M., Espinal, A., Okubo, E., and Li, B. (2009). RAP1 is essential for silencing telomeric variant surface glycoprotein genes in Trypanosoma brucei. Cell 137, 99–109. 10.1016/j.cell.2009.01.037.

18. Cestari, I., McLeland-Wieser, H., and Stuart, K. (2019). Nuclear Phosphatidylinositol 5-Phosphatase Is Essential for Allelic Exclusion of Variant Surface Glycoprotein Genes in Trypanosomes. Mol Cell Biol 39. 10.1128/MCB.00395-18.

19. Touray, A.O., Rajesh, R., Isebe, T., Sternlieb, T., Loock, M., Kutova, O., and Cestari, I. (2023). PI(3,4,5)P3 allosteric regulation of repressor activator protein 1 controls antigenic variation in trypanosomes. Elife 12. 10.7554/eLife.89331.

20. Cestari, I., and Stuart, K. (2015). Inositol phosphate pathway controls transcription of telomeric expression sites in trypanosomes. Proc Natl Acad Sci U S A 112, E2803–2812. 10.1073/pnas.1501206112.

21. Cestari, I., Haas, P., Moretti, N.S., Schenkman, S., and Stuart, K. (2016). Chemogenetic Characterization of Inositol Phosphate Metabolic Pathway Reveals Druggable Enzymes for Targeting Kinetoplastid Parasites. Cell Chem Biol 23, 608–617. 10.1016/j.chembiol.2016.03.015.

22. Kalkhof, S., and Sinz, A. (2008). Chances and pitfalls of chemical cross-linking with amine-reactive N-hydroxysuccinimide esters. Anal Bioanal Chem 392, 305–312. 10.1007/s00216-008-2231-5.

23. Merkley, E.D., Rysavy, S., Kahraman, A., Hafen, R.P., Daggett, V., and Adkins, J.N. (2014). Distance restraints from crosslinking mass spectrometry: mining a molecular dynamics simulation database to evaluate lysine-lysine distances. Protein Sci 23, 747–759. 10.1002/pro.2458.

24. Akiyoshi, B., and Gull, K. (2014). Discovery of unconventional kinetochores in kinetoplastids. Cell 156, 1247–1258. 10.1016/j.cell.2014.01.049.

25. Raschle, M., Smeenk, G., Hansen, R.K., Temu, T., Oka, Y., Hein, M.Y., Nagaraj, N., Long, D.T., Walter, J.C., Hofmann, K., et al. (2015). DNA repair. Proteomics reveals dynamic assembly of repair complexes during bypass of DNA cross-links. Science 348, 1253671. 10.1126/science.1253671.

26. Nagasaka, K., Davidson, I.F., Stocsits, R.R., Tang, W., Wutz, G., Batty, P., Panarotto, M., Litos, G., Schleiffer, A., Gerlich, D.W., and Peters, J.M. (2023). Cohesin mediates DNA loop extrusion and sister chromatid cohesion by distinct mechanisms. Mol Cell 83, 3049–3063 e3046. 10.1016/j.molcel.2023.07.024.

27. Zhou, Q., Lee, K.J., Kurasawa, Y., Hu, H., An, T., and Li, Z. (2018). Faithful chromosome segregation in Trypanosoma brucei requires a cohort of divergent spindle-associated proteins with distinct functions. Nucleic Acids Res 46, 8216–8231. 10.1093/nar/gky557.

28. Staneva, D.P., Carloni, R., Auchynnikava, T., Tong, P., Rappsilber, J., Jeyaprakash, A.A., Matthews, K.R., and Allshire, R.C. (2021). A systematic analysis of Trypanosoma brucei chromatin factors identifies novel protein interaction networks associated with sites of transcription initiation and termination. Genome Res 31, 2138–2154. 10.1101/gr.275368.121.

29. Hansen, A.S., Pustova, I., Cattoglio, C., Tjian, R., and Darzacq, X. (2017). CTCF and cohesin regulate chromatin loop stability with distinct dynamics. Elife 6. 10.7554/eLife.25776.

30. McArthur, E., and Capra, J.A. (2021). Topologically associating domain boundaries that are stable across diverse cell types are evolutionarily constrained and enriched for heritability. Am J Hum Genet 108, 269–283. 10.1016/j.ajhg.2021.01.001.

31. Ray, J., Munn, P.R., Vihervaara, A., Lewis, J.J., Ozer, A., Danko, C.G., and Lis, J.T. (2019). Chromatin conformation remains stable upon extensive transcriptional changes driven by heat shock. Proc Natl Acad Sci U S A 116, 19431–19439. 10.1073/pnas.1901244116.

32. Clayton, C. (2019). Regulation of gene expression in trypanosomatids: living with polycistronic transcription. Open Biol 9, 190072. 10.1098/rsob.190072.

33. Touray, A.O., Sternlieb, T., Isebe, T., and Cestari, I. (2023). Identifying Antigenic Switching by Clonal Cell Barcoding and Nanopore Sequencing in Trypanosoma brucei. Bio Protoc 13, e4904. 10.21769/BioProtoc.4904.

34. Cestari, I., and Stuart, K. (2020). The phosphoinositide regulatory network in Trypanosoma brucei: Implications for cell-wide regulation in eukaryotes. PLoS Negl Trop Dis 14, e0008689. 10.1371/journal.pntd.0008689.

35. DuBois, K.N., Alsford, S., Holden, J.M., Buisson, J., Swiderski, M., Bart, J.M., Ratushny, A.V., Wan, Y., Bastin, P., Barry, J.D., et al. (2012). NUP-1 Is a large coiled-coil nucleoskeletal protein in trypanosomes with lamin-like functions. PLoS Biol 10, e1001287. 10.1371/journal.pbio.1001287.

36. Soltysik, K., Ohsaki, Y., Tatematsu, T., Cheng, J., Maeda, A., Morita, S.Y., and Fujimoto, T. (2021). Nuclear lipid droplets form in the inner nuclear membrane in a seipin-independent manner. J Cell Biol 220. 10.1083/jcb.202005026.

37. (!!! INVALID CITATION !!! (2)).

38. Cestari, I. (2019). Identification of Inositol Phosphate or Phosphoinositide Interacting Proteins by Affinity Chromatography Coupled to Western Blot or Mass Spectrometry. J Vis Exp. 10.3791/59865.

39. Deutsch, E.W., Mendoza, L., Shteynberg, D., Slagel, J., Sun, Z., and Moritz, R.L.J.P.C.A. (2015). Trans-Proteomic Pipeline, a standardized data processing pipeline for large-scale reproducible proteomics informatics. 9, 745–754.

40. Hoopmann, M.R., Shteynberg, D.D., Zelter, A., Riffle, M., Lyon, A.S., Agard, D.A., Luan, Q., Nolen, B.J., MacCoss, M.J., Davis, T.N., and Moritz, R.L. (2023). Improved Analysis of Cross-Linking Mass Spectrometry Data with Kojak 2.0, Advanced by Integration into the Trans-Proteomic Pipeline. J Proteome Res 22, 647–655. 10.1021/acs.jproteome.2c00670.

41. Shannon, P., Markiel, A., Ozier, O., Baliga, N.S., Wang, J.T., Ramage, D., Amin, N., Schwikowski, B., and Ideker, T. (2003). Cytoscape: a software environment for integrated models of biomolecular interaction networks. Genome Res 13, 2498–2504. 10.1101/gr.1239303.

42. Riffle, M., Jaschob, D., Zelter, A., and Davis, T.N. (2016). ProXL (protein cross-linking database): a platform for analysis, visualization, and sharing of protein cross-linking mass spectrometry data. Journal of Proteome Research 15, 2863–2870.

43. Graham, M., Combe, C., Kolbowski, L., and Rappsilber, J. (2019). xiView: A common platform for the downstream analysis of Crosslinking Mass Spectrometry data. BioRxiv, 561829.

44. Tyanova, S., Temu, T., Sinitcyn, P., Carlson, A., Hein, M.Y., Geiger, T., Mann, M., and Cox, J. (2016). The Perseus computational platform for comprehensive analysis of (prote)omics data. Nat Methods 13, 731–740. 10.1038/nmeth.3901.

45. Zhong, J.Y., Niu, L., Lin, Z.B., Bai, X., Chen, Y., Luo, F., Hou, C., and Xiao, C.L. (2023). High-throughput Pore-C reveals the single-allele topology and cell type-specificity of 3D genome folding. Nat Commun 14, 1250. 10.1038/s41467-023-36899-x.

46. Li, H., and Durbin, R. (2010). Fast and accurate long-read alignment with Burrows-Wheeler transform. Bioinformatics 26, 589–595. 10.1093/bioinformatics/btp698.

47. Open2C, Abdennur, N., Fudenberg, G., Flyamer, I.M., Galitsyna, A.A., Goloborodko, A., Imakaev, M., and Venev, S.V. (2024). Pairtools: From sequencing data to chromosome contacts. PLoS Comput Biol 20, e1012164. 10.1371/journal.pcbi.1012164.

48. Abdennur, N., and Mirny, L.A. (2020). Cooler: scalable storage for Hi-C data and other genomically labeled arrays. Bioinformatics 36, 311–316. 10.1093/bioinformatics/btz540.

49. Kruse, K., Hug, C.B., and Vaquerizas, J.M. (2020). FAN-C: a feature-rich framework for the analysis and visualisation of chromosome conformation capture data. Genome Biol 21, 303. 10.1186/s13059-020-02215-9.

50. Wolff, J., Rabbani, L., Gilsbach, R., Richard, G., Manke, T., Backofen, R., and Gruning, B.A. (2020). Galaxy HiCExplorer 3: a web server for reproducible Hi-C, capture Hi-C and single-cell Hi-C data analysis, quality control and visualization. Nucleic Acids Res 48, W177–W184. 10.1093/nar/gkaa220.

51. Magnitov, M.D., Garaev, A.K., Tyakht, A.V., Ulianov, S.V., and Razin, S.V. (2022). Pentad: a tool for distance-dependent analysis of Hi-C interactions within and between chromatin compartments. BMC Bioinformatics 23, 116. 10.1186/s12859-022-04654-6.

