## Supplementary material for "Chromosome compartment assembly is essential for subtelomeric gene silencing in trypanosomes": Table S3

**Table S3. List of proteins that interact with PIP5Pase in *T. brucei* procyclic forms.** Proteins identified by PIP5Pase-V5 immunoprecipitation with anti-V5 antibodies followed by mass spectrometry. FC, fold-change comparing immunoprecipitated proteins from V5-tagged procyclic forms lysate to immunoprecipitations of procyclic forms lysate not expressing V5-tagged PIP5Pase. Only nuclear proteins validated by localization assays are shown.

| Gene_ID | Product Description | FC (log2) | p-value | No. peptides |
| --- | --- | --- | --- | --- |
| Tb927.11.6270 | Phosphatidylinositol phosphate 5-phosphatase | 8.94 | 5.75E-05 | 2,392 |
| Tb927.8.3950 | GGDEF domain-containing protein, putative | 4.73 | 5.92E-05 | 147 |
| Tb927.9.10770 | Polyadenylate-binding protein 2 | 4.73 | 1.04E-07 | 101 |
| Tb927.11.370 | Repressor activator protein 1 | 4.72 | 2.49E-04 | 256 |
| Tb927.11.14000 | Nuclear RNA binding domain 1 | 4.55 | 9.30E-06 | 36 |
| Tb927.3.1120 | GTP-binding nuclear protein rtb2, putative | 4.20 | 3.33E-06 | 22 |
| Tb927.1.120 | Retrotransposon hot spot protein 4 (RHS4), putative | 3.07 | 1.61E-04 | 12 |
| Tb927.5.4260 | Histone H4, putative | 2.72 | 5.48E-03 | 10 |
| Tb927.10.14750 | Fibrillarin 2 | 2.21 | 4.83E-04 | 14 |
| Tb927.7.2940 | histone H2A, putative | 2.17 | 8.51E-04 | 10 |
| Tb927.10.1060 | T-complex protein 1, delta subunit, putative | 1.89 | 1.19E-03 | 14 |
| Tb927.10.540 | ATP-dependent RNA helicase SUB2, putative | 1.82 | 5.27E-03 | 9 |
| Tb927.8.2290 | Hypothetical protein, conserved | 1.81 | 1.06E-02 | 23 |
| Tb927.1.2550 | Histone H3, putative | 1.53 | 1.57E-02 | 5 |
| Tb927.4.3060 | Hypothetical protein, conserved | 1.47 | 2.31E-02 | 5 |
| Tb927.10.2370 | Lupus La protein homolog, putative | 1.38 | 1.06E-02 | 7 |
| Tb927.9.12510 | ATP-dependent DEAD/H RNA helicase, putative | 1.38 | 2.97E-03 | 18 |
| Tb927.8.4660 | Hypothetical protein, conserved (Exportin 7 related) | 1.35 | 3.54E-02 | 8 |
| Tb927.8.3750 | Nucleolar protein 56 | 1.32 | 4.33E-02 | 19 |
| Tb927.9.5320 | Nucleolar RNA binding protein, putative | 1.23 | 5.99E-02 | 10 |
| Tb927.10.11660 | THO complex subunit 5, putative | 1.16 | 1.31E-02 | 10 |
| Tb927.10.1560 | Nucleolar protein 91 | 1.12 | 5.28E-02 | 45 |
| Tb927.9.5190 | Proliferating cell nuclear antigen | 1.06 | 2.84E-02 | 6 |
| Tb927.6.1470 | Hypothetical protein, conserved | 1.05 | 8.72E-02 | 12 |
| Tb927.9.15060 | rRNA processing protein, putative | 0.95 | 5.84E-02 | 10 |
| Tb927.10.7500 | Fibrillarin | 0.91 | 3.59E-02 | 12 |
| Tb927.8.3521 | Hypothetical protein | 0.85 | 1.89E-02 |  |
| Tb927.11.9760 | mRNA cleavage and polyadenylation factor CLP1 | 0.71 | 5.77E-02 | 12 |
| Tb927.10.15370 | DNA-directed RNA polymerases I and III subunit RPAC1, putative | 0.70 | 3.69E-02 | 4 |
