## Supplementary material for "Chromosome compartment assembly is essential for subtelomeric gene silencing in trypanosomes": Table S4

**Table S4.** **Statistics of Hi-C sequencing, alignment, and contacts.** Sequencing performed by Illumina NovaSeq. R1 and R2 refer to Illumina paired reads. * Sequencing performed with Oxford nanopore sequencer MinION. RP, biological replicates.

| **Samples** | **Group** | **Reads (R1+R2)** | **Total alignments** | **Contacts** |
| --- | --- | --- | --- | --- |
| SM427 | RP1 | 478,954,372 | 520,726,936 | 84,571,635 |
| SM427 | RP2 | 474,518,690 | 463,729,357 | 66,180,319 |
| SM427* | RP3 | 1,987,996 | 21,351,232 | 4,250,818 |
| CN PIP5Pase Tet + | RP1 | 474,518,690 | 450,581,626 | 70,610,174 |
| CN PIP5Pase Tet + | RP2 | 584,473,958 | 568,216,739 | 116,696,341 |
| CN PIP5Pase Tet + | RP3 | 432,674,790 | 472,262,320 | 111,463,674 |
| CN PIP5Pase Tet - | RP1 | 510,936,252 | 498,796,338 | 85,736,879 |
| CN PIP5Pase Tet - | RP2 | 475,100,422 | 514,872,190 | 101,158,068 |
| CN PIP5Pase Tet - | RP3 | 337,485,098 | 366,270,012 | 68,026,438 |
