## Supplementary material for "Chromosome compartment assembly is essential for subtelomeric gene silencing in trypanosomes": Table S5

**Table S5.** **Hi-C statistics of combined biological replicates.** The number of cis and trans contacts is also shown.

| Samples | Total contacts | Cis-contacts | Trans-contacts |
| --- | --- | --- | --- |
| SM427 | 155,002,772 | 117,721,249 | 37,281,523 |
| CN PIP5Pase Tet + | 298,770,189 | 264,102,431 | 34,667,758 |
| CN PIP5Pase Tet - | 254,921,385 | 210,337,093 | 44,584,292 |
