## Supplementary figures and images for "Chromosome compartment assembly is essential for subtelomeric gene silencing in trypanosomes"

### Figure S1

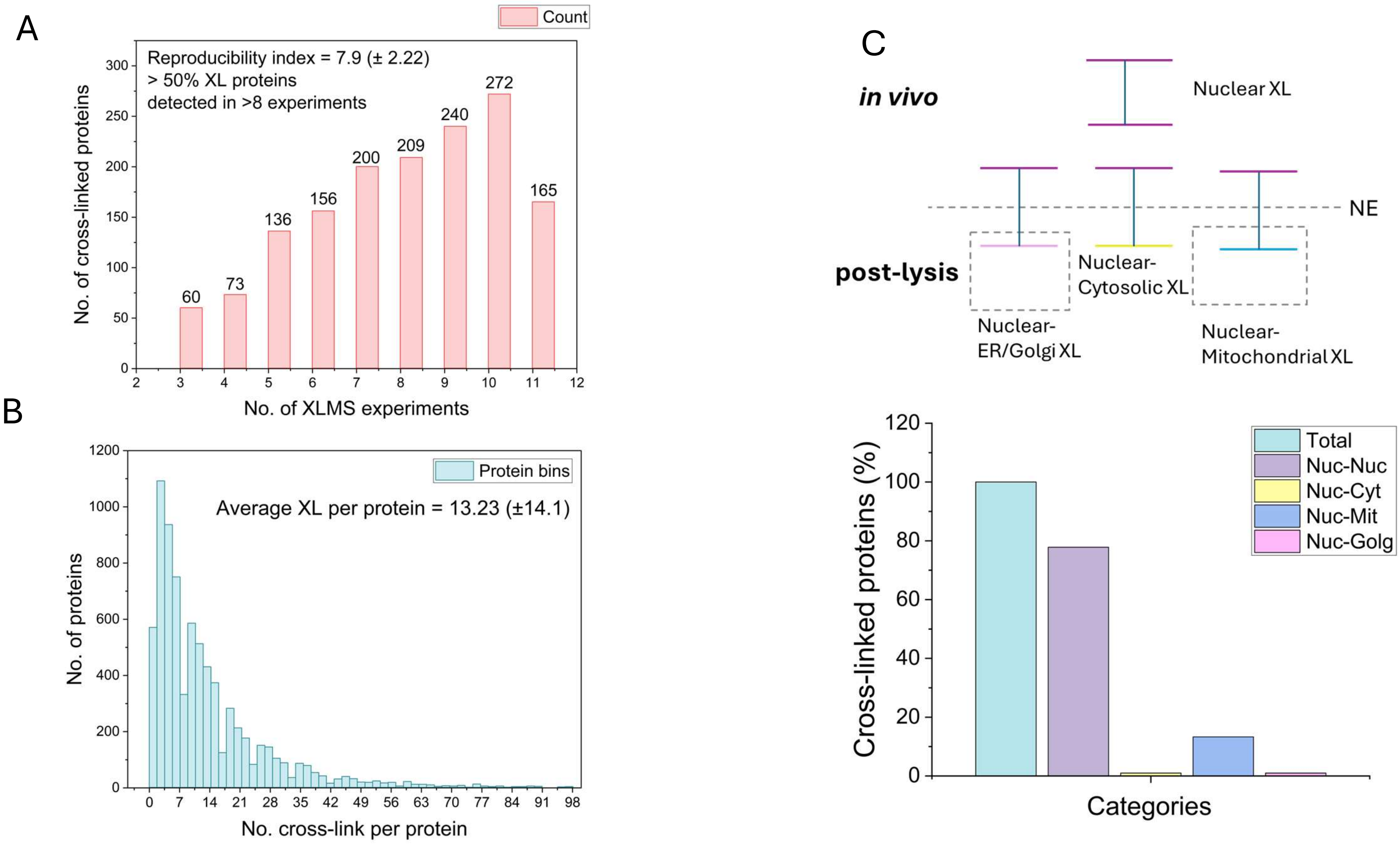

### Figure S2

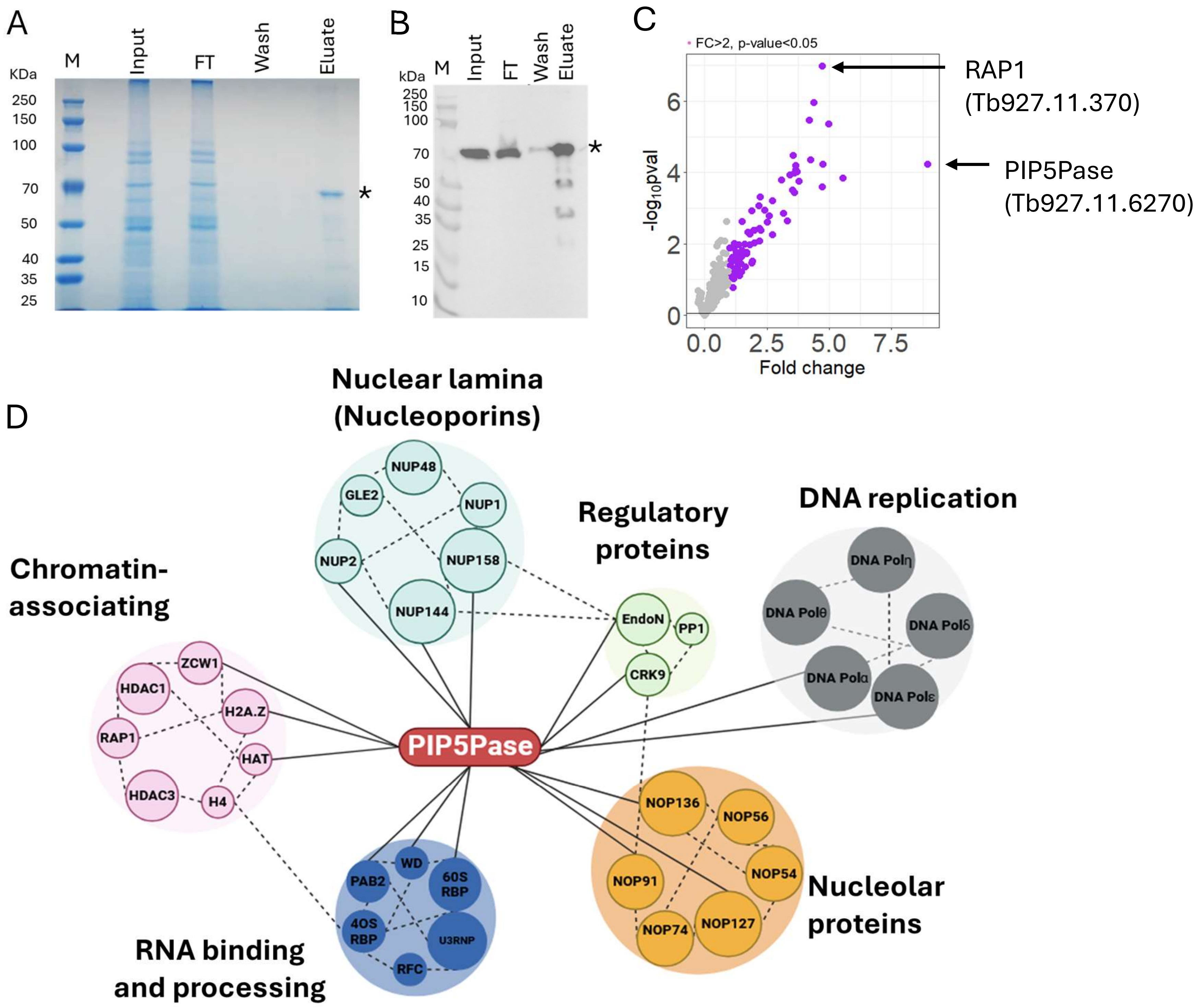
